# Selective-plane Functional Ultrasound Neuroimaging

**DOI:** 10.1101/2025.06.12.659275

**Authors:** Rick Waasdorp, Twan Gouwerok, Eleonora Munoz Ibarra, Flora Nelissen, Guillaume Renaud, Valeria Gazzola, David Maresca, Baptiste Heiles

## Abstract

Functional ultrasound (fUS) is a sensitive neuroimaging technique that uses high frame rate ultrasound to monitor brain hemodynamics as a proxy for neural activity. Recent studies have demonstrated its potential for brain-machine interfaces (BMIs) in both primates and humans. However, current 2D fUS approaches are limited to a single brain slice, restricting the ability to decode widespread neural activity. While 3D fUS could overcome this, it demands high data throughput, increased computation, and higher temperature increases due to the requirement of a higher number of transmitted waves. To address this, we present selective-plane fUS, a method that leverages the wide field of view of row-column addressed (RCA) transducer arrays to capture activity in targeted brain regions without moving the probe. By electronically selecting imaging planes, this approach achieves higher spatio-temporal resolution with lower data and pulse repetition rate compared to 3D fUS, while preserving sensitivity to neurovascular signals. Our pipeline begins with a 3D functional activation scan to guide plane selection, followed by high frame rate focused wave (FW) imaging in coronal or sagittal slices. This method offers robust detection of visually evoked responses in rodents and reduces signal variability compared to 3D fUS. By imaging only functional regions of interest, selective-plane fUS cuts computational load by an order of magnitude, enables continuous 1000 Hz recordings, and reduces functional signal variability fivefold. We envision that this method will allow tailored continuous functional imaging of widespread neuronal activity in the human brain in a BMI context.

## 1 Introduction

fUS imaging has emerged as a valuable tool for detecting neural activity in preclinical and clinical research (1, 2). Thanks to the high scalability of ultrasound, fUS has been applied to various neural systems across a wide range of species, including rodents (3–6), pigeons (7), ferrets (8), non-human primates (9, 10) and humans (11–13). Recently, fUS has garnered significant interest from the field of Brain Machine Interfaces (BMIs). BMIs enable direct communication between the brain and external devices, allowing users to control computers or prosthetics (14). By decoding the brain functions, neural implants can offer solutions for people with reduced mobility (15) or speech disabilities (16). Critical requirements for BMIs are portability and real-time imaging to allow closed-loop control, which is essential for accurate translation of the user’s neural signals to intended actions (17, 18). fUS is a good candidate for BMI due to its attractive spatio-temporal resolution (100µm and 100ms at 15 MHz), high sensitivity to slow blood flow (> 2mm/s) (19), portability, and large field of view depth-resolved imaging capabilities (20).

The first demonstration of fUS for BMI showed that it was capable of predicting movement intentions of non-human primates on a single-trial basis (21). This work relied on offline analysis of prerecorded data. Subsequent work made advances by improving the decoding ability from 2 to 8 movement directions, and allowing online, closed-loop control (22). These first implementations of fUS-BMIs were constrained to a single imaging plane, and therefore miss potentially informative functional signals out of the selected plane. They also suffer from motion of the brain during long acquisitions which means the informative functional signal could go out of the plane of imaging. To further advance fUS-BMI performance, there is growing interest in applying volumetric fUS to capture activations in a larger field of view, with the perspective of improved decoder accuracy and extension of possible actions.

Rabut *et al*. (23) and Brunner *et al*. (6) pioneered whole brain volumetric fUS using a fully populated 2D matrix array with 1024 elements. In current implementations, imaging with the full aperture requires multiplexed acquisitions (6), thereby reducing the frame rate, or connecting multiple scanners (24), decreasing portability. Moreover, due to fabrication constraints, 2D matrix imaging remains limited to a relatively small field of view with piezoelectric elements several times the wavelength, decreasing imaging quality. Addressing the full aperture also requires many channels, leading to high memory and processing bandwidth demands, incompatible with BMIs requirements. To address these limitations in the context of fUS imaging, Row Column Addressed (RCA) arrays have been proposed as a promising alternative (25, 26).

RCA transducers consist of two stacked 1D arrays with long thin line transducer elements that extend the whole aperture. The two 1D arrays (rows and columns) are oriented orthogonally to each other (Fig. 1a), allowing cylindrical focusing in transmit with one array, and cylindrical receive focusing with the other.

**Fig. 1.**
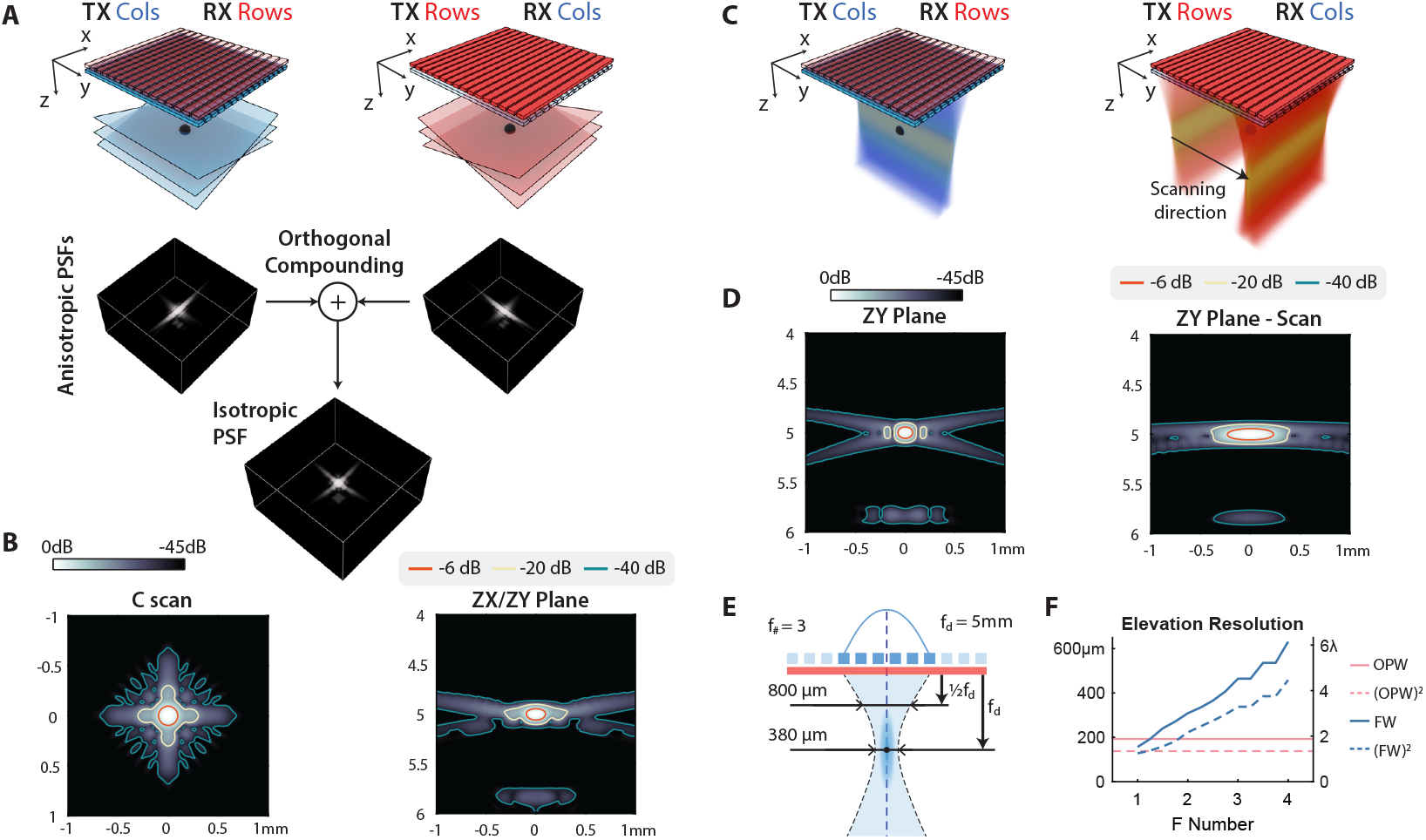
Transmission schemes for Orthogonal Plane Wave (OPW) and Focused Wave (FW) imaging simulated in Field-II. **(A)** OPW imaging of a point target with a row-column addressed (RCA) array. The sequence first transmits angled plane waves using the columns and receives with the rows (CR), after which the rows transmit and the columns receive (RC). Next, the RC and CR volumes are formed using 3D delay-and-sum beamforming and angular compounding. The RC and CR volumes have an anisotropic point spread function (PSF), and are orthogonally compounded to retrieve the final reflectional symmetric image of the medium. The shown volumetric PSFs follow from a Field-II simulation of a single point target at 5 mm depth, centered below the array. **(B)** The C-scan is shown on the left, and the B-scan is shown on the right. Due to the reflectional symmetry, the ZX and ZY planes are identical. **(C)** FW imaging of a point target with a row-column addressed (RCA) array. Transmitting a focused wave with the columns insonifies the medium with an elliptic cylinder, allowing to form a 2D image in the ZY plane.**(D)** By changing the roles of the arrays, the elliptic cylinder can insonify the orthogonal ZX plane. The position of the focus can be changed, by changing the active aperture. We simulated a plane-by-plane scan by moving the active aperture with increments of one element, which was used to form a volumetric image of the point target. The center plane of this volume was used to determine the elevation resolution. **(E)** The beam width at focus depth *f*_*d*_ = 5mm and 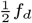 is shown bottom, for transmit *f*_#_ = 3. **(F)** The elevation resolution for different simulated transmit f-numbers (*f*_#_) for B-mode (solid lines) and the square of the B-mode (dashed lines) to find the power Doppler resolution. In **(B, D)**, the orange, yellow and blue lines indicate the −6, −20 and −40dB isolines, respectively.

Compared to a fully populated 2D matrix array, the channel count is significantly reduced from *N* ^2^ to 2*N*, eliminating the need for multiplexing and lowering memory bandwidth requirements. The RCA architecture allows production of larger aperture arrays with a lower channel count, thereby significantly increasing the field of view. RCAs have successfully demonstrated whole brain 4D fUS in rats (27), 3D anatomical imaging (28) and 3D imaging of cellular function (19). However, due to their inherent degraded reception focusing in one direction, RCA sequences to date require many transmits to reach sufficient resolution and sensitivity, decreasing frame rate and increasing the transmitted power in tissue which should be limited to guarantee safety. Although both 2D matrix and RCA arrays have significantly advanced volumetric fUS imaging, the current implementations still require substantial memory and computational bandwidth. This poses a challenge in achieving low-latency, real-time 4D fUS imaging, and hinders the application of volumetric imaging in BMIs.

There is a need for data-efficient methods to image functional brain regions with high spatio-temporal resolution while retaining an adaptive field of view. In this study we address this need, by leveraging the RCA architecture to transmit focused waves and selectively insonify functional areas of interest at high frame rates. Our working hypothesis is that task- or stimulation-related brain activity is spatially sparse. Our approach enables sampling of the brain with highly reduced memory bandwidth requirements, and simultaneously covers coronal and sagittal planes, without the need of moving the transducer. Furthermore, the plane selection is electronically reconfigurable, allowing for a flexible and adaptive wide field of view in real-time, an interesting capability in the context of permanently implanted prosthetics.

## 2 Methods

### 2.1 Simulation of Selective-plane fUS Imaging

To estimate the resolution of orthogonal plane wave (OPW) and focused wave (FW) imaging using a row-column addressed (RCA) array, we first simulated a single point scatterer in Field II (29). We modelled an RCA of 80 elements per array, 0.11 mm pitch, a center frequency of 15 MHz and 80% frequency bandwidth. We defined the medium with a speed of sound of 1540 m/s, and simulated a single point scatter at a depth of 5 mm centered below the array.

#### 2.1.1 OPW Sequence

32 plane waves with angles ranging from ± 15.5° are transmitted with the columns and then received with the rows (Fig. 1a). Using a conventional delay-and-sum beamforming algorithm, we obtain a volume presenting an elongated point spread function (PSF) in the direction of the receiving elements. For the second set of transmits, the role of the rows and columns are reversed. The beamformed data of these two orthogonal transmit sets is then summed to obtain the final volume (Fig. 1a). The resulting PSF presents a reflectional symmetry along the direction of the two orthogonal arrays leading to degraded resolution compared to fully addressed arrays. To suppress grating lobes and achieve sufficient sensitivity for functional imaging, numerous transmissions per volume are required. Finally, we determined the axial, lateral and elevation resolution in the C-scan and B-mode through the PSF using the full width at half maximum (FWHM). Furthermore, we squared the PSFs to find the power Doppler resolution (4).

#### 2.1.2 FW Sequence

A focused wave was transmitted with the rows. The resulting waves propagate and focus in an elliptic cylinder (Fig. 1c,e), from which the echoes are received by the columns. The cylinder location can be adapted by changing the active aperture on the RCA. The image from the insonified volume is reconstructed by conventional 2D delay-and-sum beamforming. To investigate the resolution in elevation, we have transmitted several focused waves in the elevation direction.

We varied the transmit f-number and aperture size to investigate the trade-off between field of view, resolution and slice thickness. For all simulations the focus depth was set to the point target depth (5 mm). For a focal depth *d* and transmit f-number *f*, the aperture size is given by *A* = ^*d*^/_*f*_. The transmit-delays for a slice positioned at *x*_*s*_ are given by,

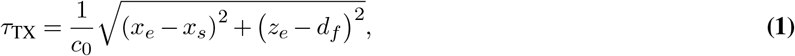

with *c*_0_ the speed of sound, *x*_*e*_ the x-coordinate of the elements, and *z*_*e*_ the z-coordinate of the elements. We transmitted *n* focused waves with the columns to scan the entire medium with increments of *p* = 0.11mm of the focal point lateral position, with *n*,

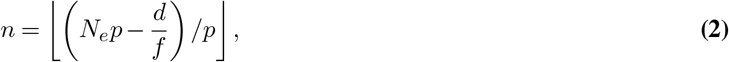

with *N*_*e*_ the number of elements and pitch *p*. The reconstructed images were concatenated, and used to determine the elevation resolution and contrast of the point target (Fig. 1d).

Finally, we determined the axial, lateral and elevation resolution in the B-mode using the full width at half maximum (FWHM). Furthermore, we squared the PSFs to find the power Doppler resolution (4).

### 2.2 Animal Procedures

All animal experiments were approved under CCD license number AVD8010020209725 at the Koninklijke Nederlandse Akademie van Wetenschappen with Study Dossier number 213601. Adult male Sprague Dawley rats weighting 470-490 g were used in this study. Animals were group housed, and kept in reversed day-night cycle with food and water available ad libitum.

#### 2.2.1 Surgery Procedure

The animal was anesthetized in a prefilled induction chamber with 5% isoflurane. After induction, the animal was transferred to a nose cone to maintain anesthesia, and placed on a heating pad of 37 °C. Anesthesia was maintained with 1-2% isoflurane, and pain was managed with carprofen (5 mg/kg) and butorphanol (2 mg/kg) with subcutaneous delivery. Dexamethasone (2.5 mg/kg)was delivered subcutaneously to prevent intracranianal edema. Eye ointment (Duratears ®) was applied to prevent ocular drying, and protective eye caps were placed over the eyes. The animal was head-fixed in a stereotactic frame and its scalp was shaved and depilated. The skin was cleaned, and a sagittal incision was made to reveal the skull after local anesthesia with Lidocaine. A 14×14 mm craniotomy was performed to reveal the brain. Saline was delivered subcutaneously to prevent dehydration. After the surgery, the animal was transferred to the imaging setup. The anesthesia was switched from isoflurane to medetomidine, initiated with a bolus of 0.5 mg/kg. After 3 minutes, we switched to continuous infusion of 0.01 mg/(kg h) delivered subcutaneously throughout the imaging session. Acoustic coupling gel was placed on the surgical window, and the RCA was positioned 2 mm above the brain and centered using live volumetric Doppler imaging.

#### 2.2.2 Visual Stimulation

To elicit brain activity, we presented visual stimuli on a 22-inch screen placed 20 cm in front of the animal (Fig. 2a). Animals were habituated to the dark prior to and in between functional recordings. Visual stimulation patterns were implemented in PsychoPy (30). We used drifting gratings (black and white) with a spatial frequency of 30°, moving at 50 °/s. The grating orientation and movement direction was switched randomly every second to multiples of 45°. The stimuli were presented for 15 seconds, followed by 30 seconds of rest. This cycle was repeated 4 times after an initial 45 seconds baseline. During rest and baseline, the screen color was gray. The total duration of one functional recording was 3m45s.

**Fig. 2.**
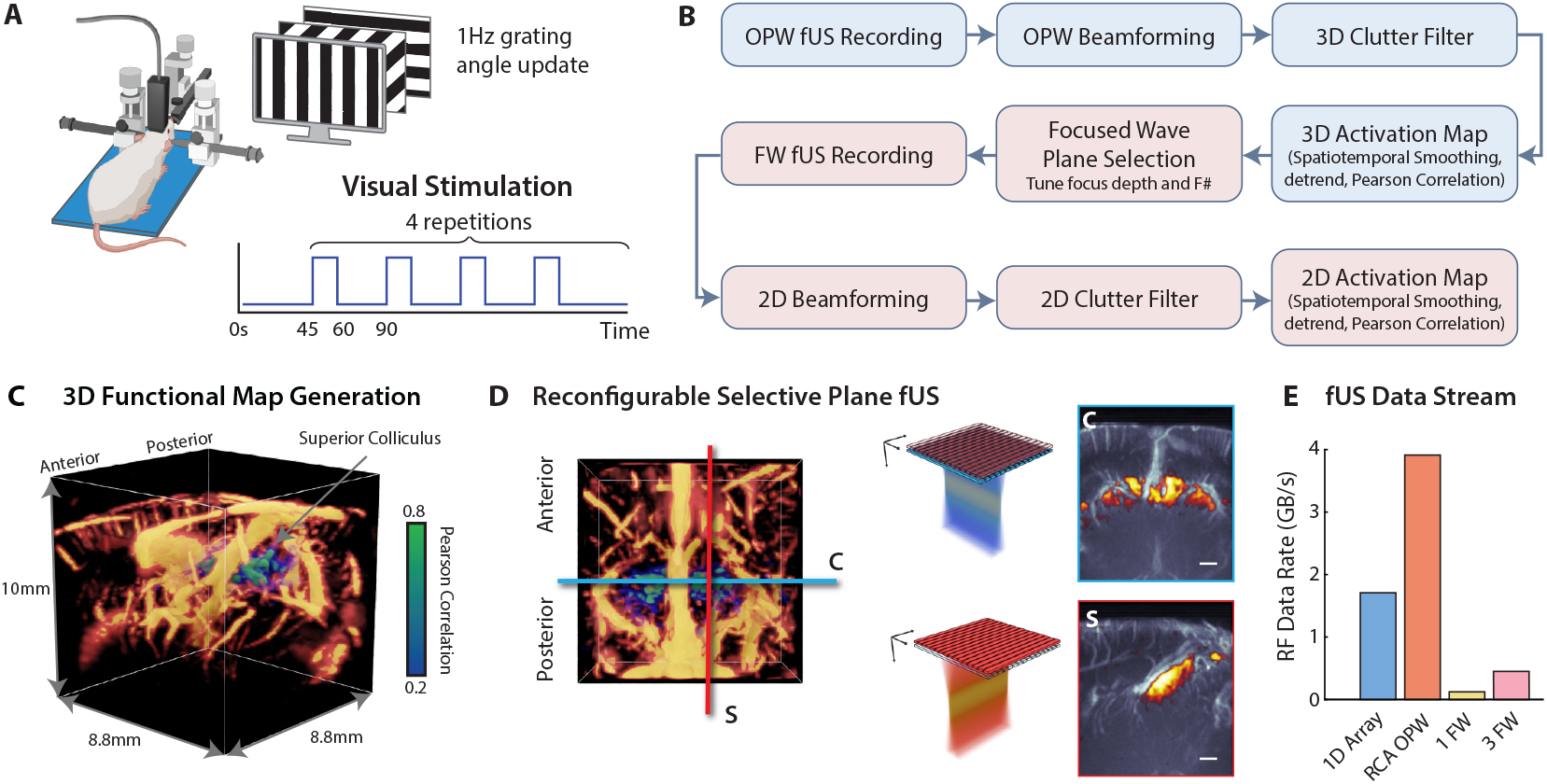
Functional selective plane imaging pipeline. **(A)** Functional imaging setup with visual stimulation. The craniotomized rat is head-fixed in a stereotactic frame, and placed on a heat pad. The row-column addressed (RCA) array is positioned above the center of the craniotomy. During functional recordings, grating bars are shown on a PC screen in 20 cm in front of the animal. The functional recording consists of an initial 45s baseline recording, followed by 4 repeats of 15s stimulation, and 30s of rest. The total recording duration is 225s. During stimulation, the grating bars on screen change orientation every 1s. **(B)** Functional data processing pipeline. **(C)** Generated 3D functional map, showing Pearson correlation overlayed on a 3D Power Doppler render. The activated region was the superior colliculus (SC). The 15MHz RCA allows to image a volume of 10×8.8×8.8 mm^3^. **(D)** Reconfigurable plane selection for the focused wave multi-plane sequence. After acquiring the 3D activation map, coronal and sagittal planes that section the activated regions can be chosen, and imaged at high frame rate. On the right example coronal and sagittal activation maps overlaid on the focused wave power Doppler are shown. **(E)** Comparison of RF data transfer rate required for 1D array imaging, RCA OPW imaging, and 1 and 3 selective planes FW imaging. Sequence parameters used to determine the data rates are listed in table 3.

#### 2.2.3 Eye Patch Experiment

To assess the stability of the functional signal under conditions of diminished neural activity, we performed an eye patch experiment. By occluding one eye with an eye patch, the activations in the contralateral hemisphere are expected to be significantly reduced, providing a controlled condition to determine the robustness of the signal.

Before conducting the eye patch experiment, we recorded 3D functional activity without eye occlusion. Based on the activity map, we selected the coronal plane for which the peak correlation was the highest, and used this plane for a selective plane functional recording. Next, we occluded the right eye, and repeated the 3D and selective plane experiments. There was a 5min rest between successive functional recordings, to avoid habituation to the visual stimuli.

### 2.3 Selective-plane Functional Imaging

Pressure waves were transmitted with an 80+80 element RCA (center frequency 15 MHz, pitch 110 µm; Verasonics, Kirkland, WA, USA) connected to a Vantage 256 channel research scanner (Verasonics, Kirkland, WA, USA). For both OPW and focused wave imaging, the imaging waveform was 2 cycles at a transmit frequency of 13.9 MHz.

#### 2.3.1 D Functional Imaging

We implemented the OPW sequence by transmitting 32 angled transmissions per array ranging from ± 15.5° with an angular pitch of 1°. After angular and orthogonal compounding, the effective volume rate was 400 Hz. We acquired Doppler ensembles of 500 ms (200 frames per ensemble), followed by a 500 ms pause, resulting in a power Doppler volume rate of 1 Hz. During acquisition, RF data was saved to disk and reconstructed in parallel in a separate thread. 3D GPU accelerated delay-and-sum beamforming was implemented in CUDA and executed on a GPU (GeForce RTX 3090, NVIDIA, USA). The resulting IQ volumes *s*_OPW_ (*x, y, z, t*) had an isotropic pixel size of 110 µm^3^, and were saved to disk for subsequent functional data processing.

We perform 3D orthogonal plane wave fUS analysis of the Power Doppler signals to create a 3D activation map (27). We elicit functional activations by presenting visual stimulation on a screen, and record the brain response with the OPW sequence. After typical spatio-temporal filtering and functional correlation (see section 2.4), we generate a 3D map of the brain activations. This map is used to select the slices for the selective-plane sequence.

#### 2.3.2 Selective-plane Sequence

After acquiring the 3D activation map, we perform selective-plane functional imaging by transmitting focused waves. The focus depth was tuned to the depth of the functional activations obtained with the 3D sequence. This was around 5 mm for the superior colliculus (SC). After simulations in Field II, we used a transmission f-number of 3, resulting in an aperture of 1.65 mm and a field of view of 7.2×8.8×10 mm^3^. We found this to be a good trade-off between elevational resolution and field of view. We used a fixed frame rate of 1000 Hz per selected plane, and ran the sequence continuously for the total duration of the functional recording. Notably, the frame rate could be increased up to the pulse repetition frequency of OPW imaging, allowing a maximum of 25.6 kHz for a single plane. Continuous recording allows overlapping of Doppler ensembles, thereby increasing the power Doppler frame rate (10). For clutter filtering, we used the same Doppler ensemble length of 500 ms as in OPW imaging, and an ensemble overlap of 400 ms, resulting in an effective power Doppler frame rate of 10 Hz.

To record visually evoked activations, we ran two subsequent recordings of the selective-plane sequence (Fig. 3a). First, we scanned the brain coronally, by selecting three planes spaced at 440 µm (4 pitch) to cover the SC, guided by the 3D activation map. Next, we recorded two sagittal planes through the left and right hemisphere, positioned at the centers of the left and right SC. The plane spacing was 2.42 mm (22 pitch).

**Fig. 3.**
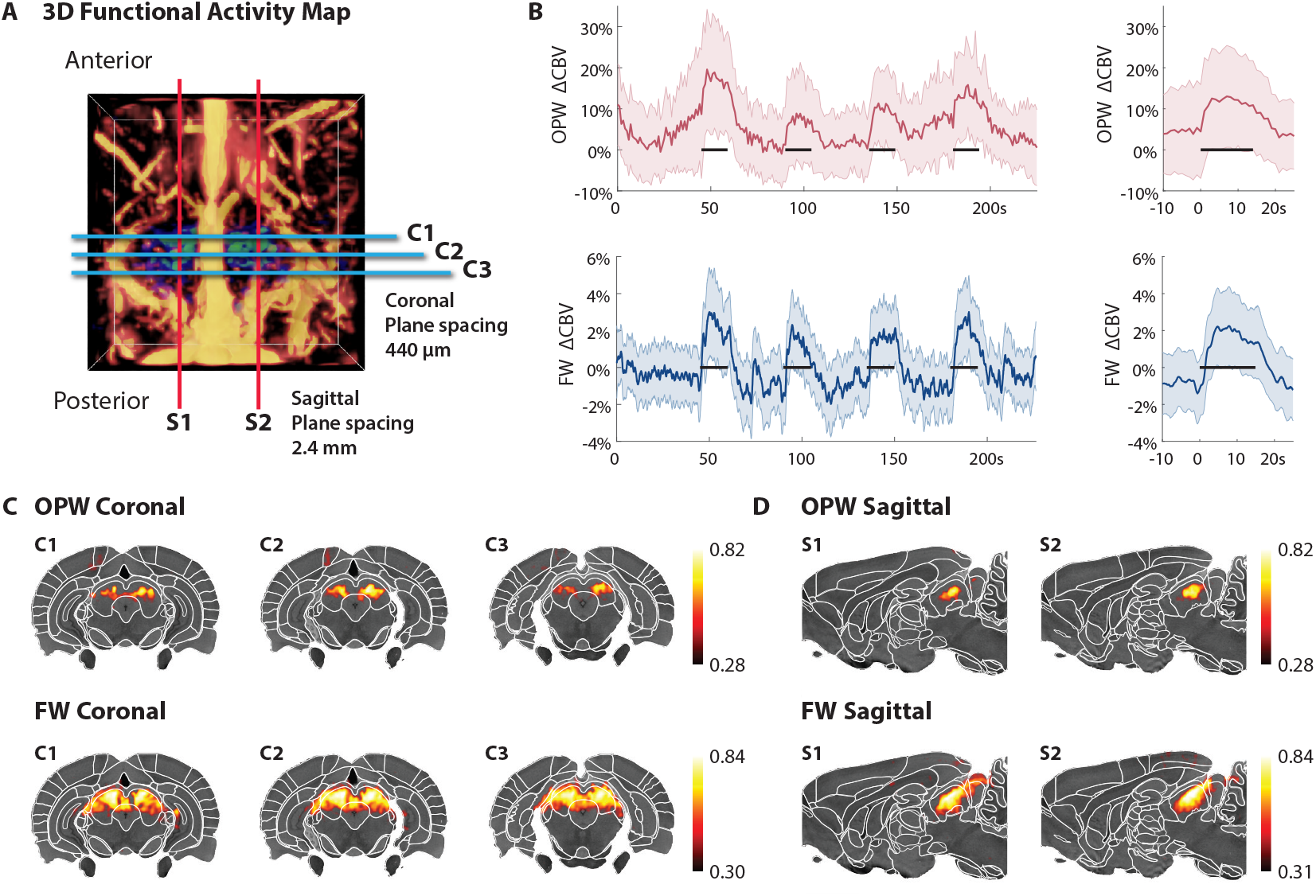
Selective-plane functional ultrasound imaging of the superior colliculus. **(A)** Orthogonal Plane Wave (OPW) generated 3D activation map, showing the selected planes for coronal and sagittal focused wave imaging (FW). **(B)** The relative increase in cerebral blood volume (CBV) during visual stimulation for OPW and FW, for significantly activated voxels. On the right, the average response for the 4 stimulation repeats is shown. **(C, D)** Neural activation overlaid on the Waxholm Space Sprague Dawley rat atlas, for OPW (top) and FW (bottom) coronal and sagittal planes. Note that the coronal and sagittal scans were acquired sequentially. FW imaging consistently reveals a larger activated brain area than OPW, and for both modalities the activation is well confined within the bounds of the Superior Colliculus. Color bars indicate the correlation threshold to the peak correlation for each recording.

Due to the fact the FW imaging requires only one transmit per region, we could run it continuously, and at higher frame rates than OPW. This significantly reduces the data collection rate, and pulse repetition frequency, as shown in Fig. 2 and table 3.

### 2.4 Functional Data Processing

#### 2.4.1 3D Activation Map Generation

After beamforming the OPW data to image volumes, blood signal was separated from static tissue using a 3D singular value decomposition (SVD) clutter filter (31). We used a fixed threshold to remove the first 50 singular vectors, since there was no motion and a minimal variation of heart rate in the anesthetized head-fixed experiments. We spatially smoothed the power Doppler volumes using a 3D Gaussian kernel with *σ* = 100 µm. Next, we temporally detrended the power Doppler signal for each pixel using a cubic polynomial (32). We determined the relative cerebral blood volume (ΔCBV) by dividing the detrended signal by the fitted trend.

The 3D functional activation map was obtained by computing the Pearson’s correlation coefficient *r*(*x, y, z*) between the normalized temporal ΔCBV signal *s*_*D*_(*t*_*i*_) and the normalized stimulus signal *a*(*t*_*i*_) using,

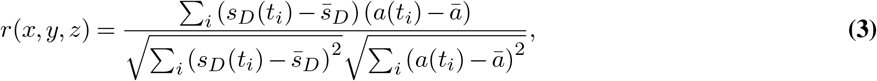

with 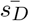 and 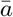 the temporal means (5). To account for the activation delay due to the neurovascular coupling, we lagged the stimulation signal by increments of one sample within a ±2s window, and computed the peak Pearson’s correlation for each lag. We finally used the correlation map of the lag that showed the maximum correlation score.

To superimpose the 3D activation map on the power Doppler volumes, we kept only voxels where the correlation score was *r > z*_score_*σ*, with *σ* is the spatial standard deviation of *r*(*x, y, z*) in a superficial noise box (10, 33). We used a *z*_score_ = 3 to find the correlation threshold *r*_OPW_.

The functional processing pipeline is shown in Fig. 2b, and a representative 3D activation map is shown in Fig. 2c.

We used the registration software developed by Brunner *et al*. (34) to manually register the volumetric OPW angiogram to the Waxholm Space atlas of the Sprague Dawley rat brain (35, 36). 3D renders were made in Blender (Blender Foundation, Amsterdam, the Netherlands) using the Scan Data Visualizer template by Cartesian Caramel (37).

#### 2.4.2 Selective-plane Functional Maps

After beamforming the selective-plane images, we performed SVD clutter filtering on each individual 2D plane. We used a fixed threshold of rejecting the first 100 singular vectors. Since frames were acquired continuously, we could compute power Doppler images using overlapping ensembles, resulting in an effective power Doppler frame rate of 10 Hz. We temporally smoothed the power Doppler signal using a 1 s moving average filter. Next, we spatially smoothed each plane with a 2D Gaussian kernel of *σ* = 100 µm, detrended using a cubic polynomial, and determined the ΔCBV by dividing the detrended signal by the fitted trend.

Finally, we computed the 2D activation maps using the Pearson’s correlation. To account for the activation delay due to the neurovascular coupling, we lagged the stimulation signal by increments of one sample within a ±2s window, and computed the peak Pearson’s correlation for each lag. We finally used the correlation map of the lag that showed the maximum correlation score. Next, we registered the selected planes using the affine transform obtained in the registration of the OPW vascular map. We used the same approach as in OPW imaging to find the correlation threshold *r*_FW_ to superimpose the activation maps on the atlas or power Doppler images. The complete functional processing pipeline is shown in Fig. 2b. Once the 3D activation map was obtained, we switched to our selective-plane functional sequence. Fig. 2d shows the top view of the 3D activation map, with two representative coronal and sagittal focused wave planes. The activity map is overlaid on the FW power Doppler for pixels above the correlation threshold.

#### 2.4.3 OPW and FW comparison

To compare the functional activation maps generated by both imaging modalities, we computed the maximum intensity projection of the 3D vascular and correlation map over an elevation range of 330 µm, or 3 pitch. This range corresponded to the FW power Doppler elevation resolution found in simulation (Fig. 1f).

To compare the CBV response of OPW and FW, we converted the OPW Doppler volumes to a set of selected planes, corresponding to the FW plane locations. For a FW plane position, we selected the corresponding OPW plane, and took the maximum intensity projection of the power Doppler maps over an elevation width of 330 µm.

## 3 Results

### 3.1 Simulation OPW and FW

We compared the PSFs of OPW and FW obtained with simulation in Fig. 1. For the OPW B-mode, we found a lateral and elevation resolution of 193 µm (1.7*λ*), and an axial resolution of 114 µm (1.0*λ*). To estimate the power Doppler resolution, we squared the PSFs, and found a lateral resolution of 137 µm (1.2*λ*) and an axial resolution of 81 µm (0.7*λ*).

For focused wave imaging, we tested different transmit f-numbers. The B-mode axial and lateral resolution were scarcely affected by the transmit f-number, and were 113±0.2 µm (1.0*λ*) and 148±0.3 µm (1.3*λ*) respectively. In the squared PSFs the axial and lateral resolution were 80±0.2 µm (0.7*λ*) and 106±0.2 µm (1.0*λ*), respectively. The elevation resolution varied with the transmit f-number, as shown in Fig. 1f. The aperture size is defined by the choice of transmit f-number and focus depth, with an aperture of 5.2 mm for *f*_#_ = 1 and 1.2 mm for *f*_#_ = 4. Given the limited available aperture of the transducer, imaging with *f*_#_ = 1 restricts the possible field of view to a 3.6×8.8×10 mm^3^ volume. To retain a wide field of view, we opted for a middle ground in elevation resolution and aperture size. For *f*_#_ = 3, the aperture size is 1.65 mm, allowing scanning of a 7.2×8.8×10 mm^3^ volume, with a power Doppler elevation resolution of 303 µm at focus. Therefore, *f*_#_ = 3 was chosen for all focused wave functional recordings.

### 3.2 3D Functional Imaging

To guide the plane selection in our selective-plane imaging, we first acquired a 3D activation map. Fig. 2c shows the activation map superimposed on the power Doppler resulting from visual stimulation. After defining a noise box, we determined the significant correlation threshold for OPW of *r*_OPW_ = 0.281. The shown volumetric correlation map only shows voxels that are above the correlation threshold. The peak correlation score was 0.817. The shape and location of the activated area corresponds well with the Superior Colliculus, as reported earlier by (3).

Fig. 3a shows the selected slices for the coronal and sagittal selective-plane recordings. For the coronal planes, we determined the maximum intensity projection of the OPW Doppler planes over 330 µm corresponding to the simulated FW elevation resolution, and spatially averaged the CBV response over all significantly activated voxels (see Fig. 3b). From this it becomes clear that the OPW CBV increased by 6.5 ± 11.9% during stimulation.

### 3.3 Selective-plane Functional Imaging

After obtaining the 3D activity map, we switched to selective-plane focused wave imaging. Fig. 3a shows the selected slices for the coronal and sagittal recordings. We spatially averaged the CBV response over all significantly activated voxels in the coronal planes (see Fig. 3b), and determined the average CBV increase across stimulation trials. The FW CBV increased by 2.3 ± 2.1% during stimulation.

Fig. 3c presents a comparison between the OPW and FW activation maps of the selected planes after registration with the Waxholm Space atlas. It becomes clear that for OPW, the activated area is confined within the borders of the SC, in both coronal and sagittal recordings. In FW, the activated area is larger than OPW, but is still confined within the borders of the SC. The correlation threshold for OPW and FW was similar, at 0.281 and 0.296 respectively. The peak correlation was 0.817 for OPW and 0.841 for FW, indicating a small increase in FW.

### 3.4 Diminished neural activity fUS stability

To compare the performance of OPW and FW under diminished neural activity, we performed an eye patch experiment. Fig. 4a,d shows the top and 3D view of the OPW activation with and without eye patch. Without eye patch, both hemispheres show activation, with similar peak correlation (difference *<* 0.05) in both hemispheres for both OPW and FW (see table 2). With the right eye patched, the left hemisphere shows lower activation, with a 0.24 decrease in correlation for OPW, and a decrease of 0.14 for FW.

**Table 1.** Average per hemisphere CBV increase during stimulation with respect to a 10-second window before stimulation, in functional recordings with and without eye patch. Values report mean ± standard deviation.

| Mode | Vision impairment | ΔCBV Left (%) | ΔCBV Right (%) |
| --- | --- | --- | --- |
| OPW | None | $28.1 \pm 16.1$ | $18.1 \pm 11.7$ |
| OPW | Left | $10.5 \pm 11.1$ | $14.9 \pm 12.8$ |
| FW | None | $11.3 \pm 4.6$ | $10.0 \pm 4.3$ |
| FW | Left | $3.2 \pm 2.0$ | $5.8 \pm 3.1$ |

**Table 2.**
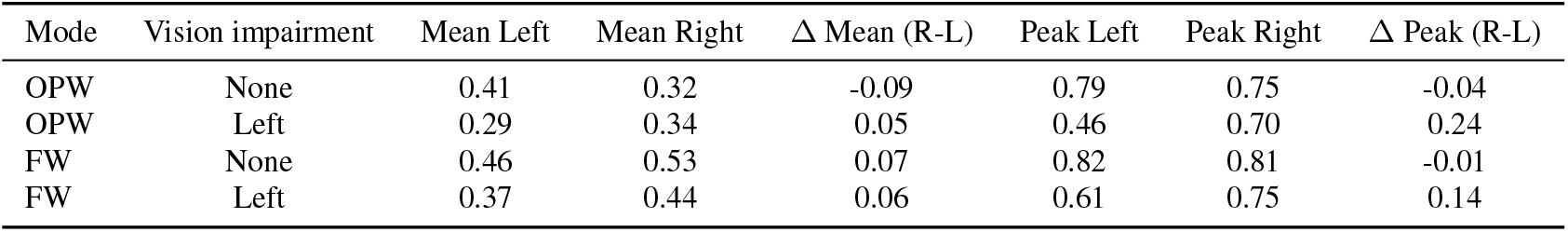
Average and peak Pearson correlation scores per hemisphere in functional OPW and FW recordings with and without eye patch. (R-L) denotes the difference between the right and left hemisphere for mean or peak correlation.

**Table 3.** Comparison of PRF and RF data rate for various imaging modes during continuous recording, assuming 200% bandwidth sampling and 10 mm deep imaging.

| Array | Imaging mode | TX per frame | Frame rate (Hz) | PRF (Hz) | Data rate (GB/s) | Ref |
| --- | --- | --- | --- | --- | --- | --- |
| 1D array | PW | 7 | 1000 | 7000 | 1.7 | (33) |
| RCA | OPW | 64 | 400 | 25600 | 3.9 | Current Study |
| RCA | 1 FW | 1 | 1000 | 1000 | 0.15 | Current Study |
| RCA | 3 FW | 3 | 1000 | 3000 | 0.46 | Current Study |

**Fig. 4.**
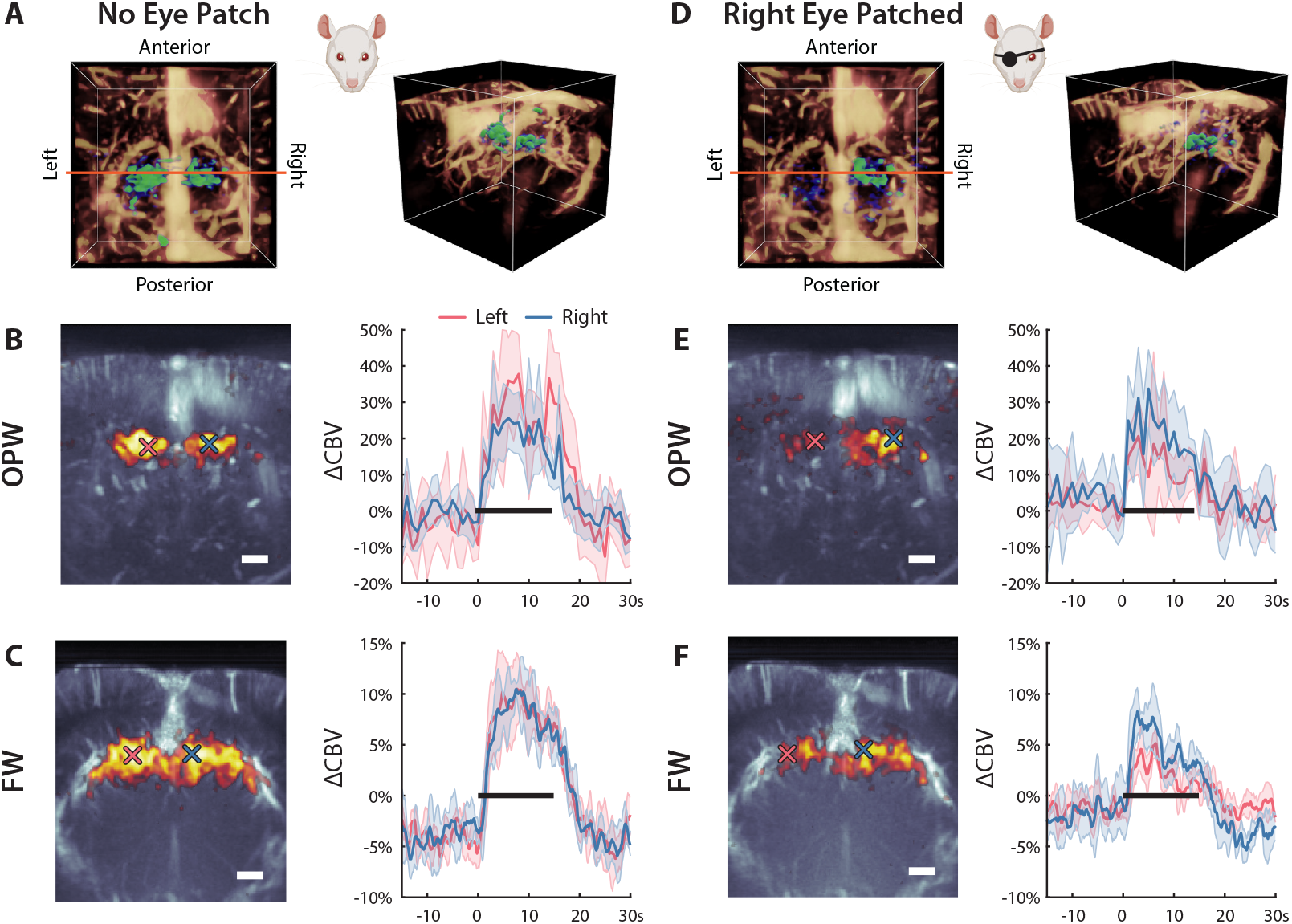
Functional ultrasound imaging of the superior colliculus with eye patch. **(A, B, C)** No eye patch. **(A)** the 3D activation map shows bilateral activation of the superior colliculus. The orange line overlaid on the top view shows the location of the focused wave imaging plane. **(B)** The OPW Pearson correlation is superimposed on the OPW power Doppler map. The blue and red crosses denote the location with maximum correlation in the left and right hemisphere, respectively. The relative cerebral blood volume (CBV) traces on the right show the hemodynamic response for the left and right hemisphere, corresponding to the pixel indicated by the crosses. **(C)** shows the activation map and relative CBV change for focused wave imaging. Both OPW and FW show a similar response in the left and right hemisphere. Although the CBV change for FW is lower than OPW, FW shows lower signal variability. **(D, E, F)** Experiment with right eye patched. Consequently, the left hemisphere shows a diminished response, resulting in lower correlation scores and lower relative CBV changes. This finding is consistent for OPW **(E)** and FW imaging **(F)**. The single pixel CBV traces for FW show lower variability in the left hemisphere compared to OPW.

Fig. 4b,e show the maximum intensity projected OPW correlation map over 330 µm superimposed on the OPW vascular map. The location of peak correlation corresponded well between the OPW and FW recordings without eye patch. In the eye patch experiments, the peak correlation location varied more, as can be seen by the crosses in Fig. 4e,f.

To determine the stability of the CBV signal in OPW and FW, we took the CBV time trace of the peak activated voxel in the left and right hemisphere, and averaged the CBV signal across stimulation repetitions. The single voxel based CBV traces shown in Fig. 4c,f showcase the 10-fold higher temporal sampling in FW than OPW. In the FW recording with eye patch, the diminished response in the left hemisphere is much more visible in the FW single voxel traces.

Table 1 presents a quantification of the average CBV increase during stimulation with respect to a 10-second window before stimulation. For both OPW and FW, the CBV response decreases in both hemispheres when one eye is patched. Furthermore, the CBV signals in the left hemisphere show lower variability, reduced from 11.1% in OPW to 2.0% in FW.

## Discussion

### Summary Key Findings

fUS has emerged as a transformative neuroimaging modality, unlocking new possibilities for both fundamental neuroscience research and a variety of (pre)clinical applications. However, current implementations of 4D fUS demand high memory throughput and computational bandwidth, presenting a key bottleneck for achieving low-latency, real-time 4D fUS imaging, crucial for clinical application such as BMIs. While future advances in computing hardware will alleviate these limitations, we present an intermediate solution that allows efficient sampling of selective-planes in the brain, relying on the hypothesis that task- or stimulation-related brain activity is spatially sparse. We realized this selective-plane imaging by transmitting focused waves with the RCA.

By transmitting a focused wave with either the rows or columns, we insonify selected brain regions with an elliptic cylinder, from which the backscatter pressure waves are received with the orthogonal array. Utilizing this partial transmit focus, followed by 2D delay-and-sum beamforming to focus in reception, we demonstrate that you only require a single transmit per plane, in contrast to the 64 transmits needed per volume in orthogonal plane wave (OPW) imaging. This reduces the RF data bandwidth by a factor 26, from 3.9 GB/s in OPW imaging to 0.15 GB/s per FW plane. Furthermore, reconstructing FW planes is computationally more efficient, as it is a single 2D beamforming procedure, opposed to the two stage orthogonal beamforming done in OPW. Conventional focused wave imaging on 1D arrays is slow due to the line by line scanning, and is not suitable for functional imaging. The RCA architecture allows focused imaging at high frame rates by turning this into plane-by-plane scanning. The choice of planes to sample was guided by registration of a 3D activation map. For this, we used functional OPW (27). After recording this 3D activation map once, the map can be reused to guide the choice of selective-planes and select optimal transmit parameters. Future experiments could instead register the power Doppler of the whole brain with an atlas to guide the selection of planes and transmit parameters.

In our simulations, we found that the FW elevation resolution is highly dependent on the transmit f-number, and was 303 µm at focus for power Doppler imaging with *f*_#_ = 3. As expected, the power Doppler elevation resolution for OPW is higher than FW, and as a result, FW integrates the vascular signal over a larger slice width. Although FW elevation resolution was higher than the OPW elevation resolution, it is comparable to the elevation resolution of a 1D array. Using a 15 MHz 1D array, Macé *et al*. (4) found a power Doppler elevation resolution of 303 µm when scanning a 20 µm wire submerged in a water tank (see supplementary figure S2 of (4)). However, in focused wave imaging with the RCA, it is important to note that the elevation resolution degrades rapidly when imaging away from the focus depth due to the absence of a focusing lens.

It is difficult to pinpoint why the OPW and FW show different activation maps and different increases in CBV during activation, as multiple implementation differences are at play. One reason is that in clutter filtering, the SVD for OPW works on a 3D volume, which results in a better separation of static tissue and blood flow compared to its 2D SVD counterpart in FW imaging, which potentially results in higher CBV changes for OPW. In FW imaging, the lower CBV changes could be explained by the reduced number of transmits per power Doppler frame, which was 500 for FW, and 12800 for OPW. The lower variability of the FW CBV signal likely is caused by the continuous sampling of the FW sequence, allowing overlap of Doppler ensembles and an increased power Doppler frame rate. To match the power Doppler rate of OPW imaging, we applied a temporal moving average filter with a 1-second window, which may contribute to the reduced CBV variability. Depending on the application, the FW frame rate and degree of temporal smoothing can be tuned to detect propagating brain activations, as demonstrated by (10).

When comparing the activation maps of OPW to FW, we see that FW reveals a larger activated area, likely due to the increased slice thickness resulting from the elevation resolution. The increase in relative CBV during stimulation was lower in FW than OPW. Although this could indicate a decreased sensitivity, the lower signal variability on individual voxel basis and comparable peak correlation scores show that FW imaging has sufficient sensitivity to detect functional activations. These results are promising for application of focused wave imaging in a fUS-BMI, where low signal variability is key to facilitate decoding of brain activity on a single trial basis.

Comparing the binocular and

### 4.2 Comparison to Existing Methods

Alternative approaches to improve brain coverage include the recently developed 1D multi-array that enables simultaneously sampling of 4 planes at a fixed distance of 2.1 mm (32), or optimized motorized sweeping to enable 3D fUS with a 1D array (38, 39). While these approaches leverage the superior image quality of a 1D array, the requirement of a motorized linear stage makes these solutions impractical for a portable BMI or for human fUS as a whole. Furthermore, they do not allow simultaneous high frame rate readout across the brain.

OPW presents an effective middle ground, allowing large field of view brain imaging at reduced channel count compared to fully populated 2D arrays. With its volumetric capabilities, it eliminates the need for motorized scanning. However, the computational requirements are high due to the large number of transmits required to achieve sufficient Doppler dynamic range (27). The large number of transmits limits achievable volume rates and leads to a very high pulse repetition frequency (see table 3), which in turn raises concerns about element overheating. In our implementation, we should have changed the angular step to 0.5 deg as recommended by Sauvage *et al*. for a more effective suppression of grating lobes (27). This makes it currently challenging to run continuous, high frame rate functional OPW recordings.

Our selective-plane imaging approach, allows to sample coronal and sagittal planes in the same functional recording, although in this study these recordings were performed subsequently. This is a benefit of using the RCA, and not possible with a traditional 1D array. Finally, aligning a 1D transducer with the correct plane covering brain regions of interest is challenging in the sagittal direction, due to the absence of distinct, symmetrical, anatomical landmarks. Here, registration of the 3D angiogram of activation map solves this issue, and allows accurate brain region alignment in both the sagittal and coronal planes.

### 4.3 Implications and Potential Applications

Our approach of selective-plane imaging is fully adaptive, and offers several potentials for fUS-BMI. Current fUS-BMI implementations rely on 2D imaging, thereby covering a limited region of the brain. Therefore, 2D imaging is unable to capture task related activations occurring outside the plane. As a result, there is growing interest in leveraging volumetric fUS to expand brain coverage, with the potential of enhanced decoder accuracy and broaden the range of possible outputs of a BMI. Current 4D fUS implementations using either fully populated 2D matrix arrays or row column addressed (RCA) arrays generate large datasets, imposing substantial demands on memory and computational bandwidth. This makes it challenging to apply volumetric imaging to a real time closed loop BMI.

Currently, fUS-BMI positions a 1D array in predefined slots in a 3D printed holder. Initially aligning these slots with the correct plane can be cumbersome, and over time, these predefined slots can misalign with the plane of interest, leading to decreased decoder performance (40). Our approach enables periodic registration of a 3D angiogram or functional map and correction via electronic aperture reconfiguration to maintain proper alignment, thereby reducing the need for precise transducer positioning. In the context of implanted BMIs, the ability to electronically steer the imaging plane offers a tremendous advantage.

BMI research has demonstrated that movement-related spatial representations are typically distributed across different brain regions (22). Therefore, simultaneously recording from multiple regions, at reduced memory and computational overhead compared to volumetric imaging, provides a significant advantage.

Although fUS-BMI research has indicated that decoding accuracy decreases with lower spatial resolution, our focused wave imaging approach allows imaging with the same spatial resolution as used by Norman *et al*. (21). Non-human primates BMI research has shown that the most informative voxels are spread across a 5 mm depth range in the cortex (22). Due to the poor elevation resolution away from focus, multiple focus depths are needed to ensure sufficient elevation resolution across depth, increasing the PRF and data rate. To mitigate increased memory and computational bandwidth requirements, the sequence can be further optimized to selectively receive only RF data around a small depth window, tuned per focus depth transmit. This highlights the adaptability of the method.

This study, together with the recently introduced nonlinear sound sheet microscopy (19), synthetic aperture imaging (41), and advances in orthogonal plane wave imaging (27, 42), demonstrate the versatility of the row column addressed transducer architecture in various applications. Together with sound-sheet microscopy, our work signals a shift in strategy: from brute-force volumetric plane wave insonification to efficient, application specific acoustic beams.

## 5 Conclusion

This study introduces selective-plane fUS, a reconfigurable ultrasound neuroimaging method. By efficiently sampling targeted brain sections, our approach significantly reduces data bandwidth and computational demands while maintaining a spatial resolution comparable to 2D fUS. Using selective-plane fUS, we successfully revealed activations in the superior colliculus in response to visual stimulation, observed similar peak correlation to 3D fUS, and measured a reduced functional signal variability. Together, our findings makes real-time, multi-region decoding of the brain possible, thereby addressing a pressing need for compact and human implantable fUS-BMI paradigms.

## 6 Acknowledgements

The authors thank V. Galligioni, G. de Fluiter, and veterinary staff of the Netherlands Institute of Neuroscience for their continuous support.

This project was partially funded through grants from the Medical Delta Ultra HB program (RW), the Chan Zuckerberg Initiative (Dynamic RFA number 2023-321233, EMI), the 4TU Precision Medicine Program (DM) and the European Union (Marie-Sklodowska Curie Fellowship MIC-101032769, BH).

